# Emergence of glycogen synthase kinase-3 interaction domain enhances phosphorylation of SARS-CoV-2 nucleocapsid protein

**DOI:** 10.1101/2022.01.24.477037

**Authors:** Jun Seop Yun, Nam Hee Kim, Hyeeun Song, So Young Cha, Kyu Ho Hwang, Jae Eun Lee, Cheol-Hee Jeong, Sang Hyun Song, Seonghun Kim, Eunae Sandra Cho, Hyun Sil Kim, Jong In Yook

## Abstract

A structural protein of SARS-CoV-2, nucleocapsid (N) protein is abundantly expressed during viral replication. The N protein is phosphorylated by glycogen synthase kinase (GSK)-3 on the serine/arginine (SR) rich motif located in disordered regions. Although phosphorylation by GSK-3β constitutes a critical event for viral replication, the molecular mechanism underlying N phosphorylation is not well understood. In this study, we found the putative alpha-helix L/FxxxL/AxxRL motif known as the GSK-3 interacting domain (GID), commonly found in many endogenous GSK-3β binding proteins, such as Axins, FRATs, WWOX and GSKIP. Indeed, N interacts with GSK-3β similarly to Axin, and Leu to Glu substitution of the GID abolished the interaction, with loss of N phosphorylation. Unlike with endogenous GID proteins, the N interaction neither disturbs endogenous GSK-3 activity nor regulates subsequent canonical Wnt activity and the Snail-EMT program. Notably, N abundance in SARS-CoV-2 is incomparably high compared to other coronaviruses, such as 229E, OC43 and HKU1. Compared to other coronaviruses, N harbors a CDK1 primed phosphorylation site and Gly-rich linker for enhanced phosphorylation by GSK-3β. Furthermore, we found that the S202R mutant found in Delta and R203K/G204R mutant found in the Omicron variant allows increased abundance and hyper-phosphorylation of N. Our observations suggest that the emergence of GID and mutations for increased phosphorylation in N may have contributed to the emergence of SARS-CoV-2 and evolution of variants, respectively. Further study, especially in a BSL3-equipped facility, is required to elucidate the functional importance of GID and N phosphorylation in SARS-CoV-2 and variants.

---

The N protein is the most abundantly expressed structural protein during viral replication of SARS-CoV-2 (1). In addition to its structural role in the ribonucleoprotein complex in virion, the N protein plays key roles in viral RNA and protein synthesis, packaging, and envelope formation (2,3). Although the spike protein is only used as an immunogen in current vaccines, serological antibodies against N protein can be used for detection of early and previous infection (4,5). The highly basic N protein of coronavirus consists of about 400 amino acids (∼ 50kDa) with three distinct domains. Structured N-terminal and C-terminal domains play diverse roles in multimerization and RNA binding (3). The central disordered region harbors the Ser/Arg (SR)-rich motif, which is highly conserved in other coronaviruses, such as OC43, HKU1 and MERS. Interestingly, the SR-rich motif of coronavirus is phosphorylated by GSK-3, and GSK-3 inhibitors suppress viral replication in Vero cells (6,7). GSK-3 inhibition also selectively reduces genomic RNA and long subgenomic mRNA (8), and a clinical trial with a GSK-3 inhibitor, lithium, yielded reduced risk of SARS-CoV2 (9). Crucially, a recent study revealed that N mutations around the SR-rich motif found in SARS-CoV-2 variants allows significantly increased viral replication compared with ancestral Wuhan Hu-1 N protein (10). These results indicate the importance of N phosphorylation by GSK-3 during viral replication and mutational evolution of coronavirus although the underlying molecular mechanism is unclear.

GSK-3 is an endogenously abundant kinase which plays a key role in many signaling pathways, including Wnt signaling. In the Wnt pathway, GSK-3 is recruited to a multi-protein complex with adenomatous polyposis coli (APC) and scaffolding protein Axin (11,12). Axin directly binds to GSK-3, resulting in inhibition of the kinase activity and regulation of canonical Wnt activity (13). Crystallography of GSK-3 and Axin peptide reveals that hydrophobic helical ridges formed by Axin residues of Phe388, Leu392, Leu396 in GID pack into the helical groove of the 285-299 loop in GSK-3 (14). This interacting domain is also found in other GSK-3 interacting proteins, such as FRATs, GSK-3 interacting protein (GSKIP), and WWOX (15,16). The GID in many proteins provides diverse functions for GSK-3 and its substrate. For example, multi-protein complex of APC and GSK-3 enhances β-catenin phosphorylation and Axin-GSK3 binding provides nuclear export of GSK-3 stabilizing, epithelial-mesenchymal transition (EMT)-inducer Snail in cancer (14,17). The typical amino acids in helical GID consist of L/FxxxL/AxxRL (Fig. 1A). In this study, we found GID located next to the SR-rich domain of N protein in SARS-CoV-2 (hereafter N), critical for N protein phosphorylation, suggesting that interaction between N and GSK-3 plays a critical role in viral replication and evolution of SARS-CoV-2. Compared to N protein in other non-pathogenic coronaviruses, N protein of SARS-CoV-2 also harbors a CDK1 phosphorylation site and a flexible linker between GID and the SR-rich region, allowing enhanced phosphorylation of N. Further, we found that N mutations in Delta and Omicron variants provide enhanced phosphorylation and increased abundance of N.

**Figure 1.**
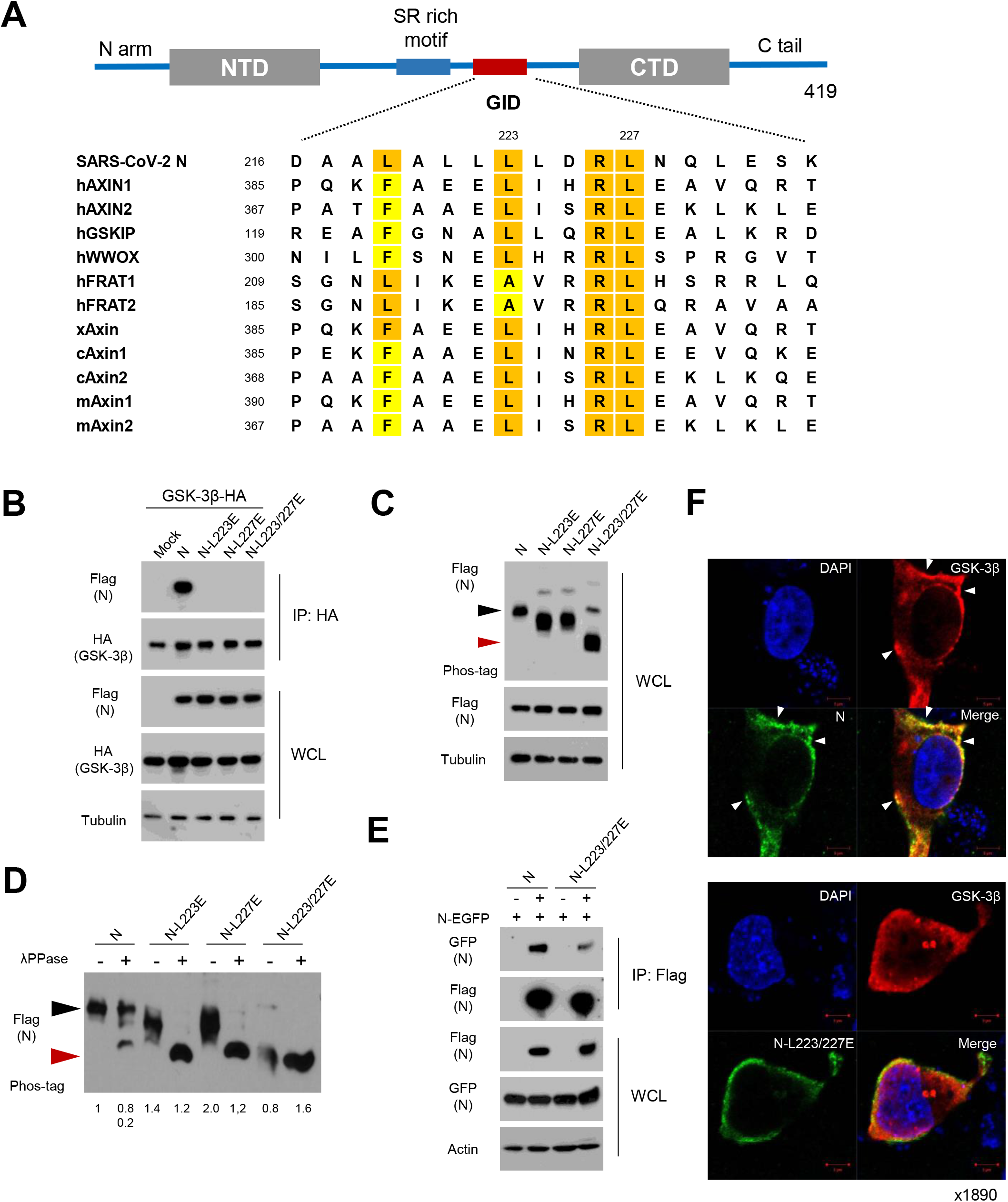
Potential GID in N of SARS-CoV-2 is responsible for phosphorylation of N. A, Schematic diagram of structural organization of SARS-CoV-2 N protein (upper). SR-rich motif and potential GID are located in the central disordered region. Sequence alignment of GID (L/FxxxL/AxxRL) in SARS-CoV-2 N and endogenous GSK-3 interacting proteins with human (h), mouse (m), xenopus (x), and chicken (c). Highlights are shown for conserved hydrophobic residues of Lue, Phe, and Ala (lower). B, Flag-tagged N and mutants harboring Lue to Gly substitutions with HA-tagged GSK-3β were expressed in 293 cells. Interaction between N or mutants and GSK-3β were determined following immunoprecipitation (IP) with anti-HA beads. Whole cell lysate (WCL) serves as input abundance for IP. C, Ancestral or mutant N expression vectors were transfected into 293 cells, and the lysates were subjected to western blot and Phos-tag gel analysis to determine protein abundance and mobility shift by phosphorylation status of N, respectively. Black and red arrowheads correspond to the fully phosphorylated and dephosphorylated N on a Phos-tag gel, respectively. D, N or mutant N lysates were subjected to lambda protein phosphatase (λPPase) followed by Phos-tag gel analysis. E, Confocal images of ancestral N or 223/227 mutant (green) and GSK-3β (red) in 293 cells. Arrows indicate co-localized foci of N and GSK-3β in condensate-like structures. Nuclear staining with DAPI and (blue); scale bar, 5 μm.

## Results

### N of SARS-CoV-2 harbors a GSK-3 interacting domain similar to AXIN

Phosphorylation of many endogenous and viral proteins by GSK-3 is dependent on GSK-3 binding of protein-protein interaction (14,18). In turn, GSK-3 interacting domain (GID) embedded in GSK-3 substrate provides efficient self-phosphorylation. Because N of coronavirus is phosphorylated by GSK-3 (6,7), we examined whether GID provides N-GSK-3 interaction and efficient phosphorylation, finding a L/FxxxLxxRL motif in the unstructured disordered region of ancestral Wuhan Hu-1 N protein (Fig. 1A). The amino acid sequence of GID is highly conserved in many GSK-3 interacting proteins, such as AXINs, GSKIP and WWOX. Because the hydrophobic residues in GID play a critical role in GSK-3 interaction (14,19,20), we generated Lue to Glu substitution mutants (L223/227E) from ancestral SARS-CoV-2 N protein. We next transfected them with HA-tagged GSK-3β (hereafter GSK-3) and N expression vectors in 293 cells and performed immunoprecipitation assay to examine interaction between N and GSK-3. Indeed, N protein interacts with GSK-3 and point mutant of hydrophobic residues in N-GID, largely abolishing the interaction with GSK-3, although protein abundance of N and mutants was comparable (Fig. 1B). To further determine the role of N-GSK-3 interaction, we next examined the phosphorylation status of N and mutants by means of mobility shift on a Phos-tag gel. Interestingly, mutation of hydrophobic residue(s) largely decreased phosphorylation of N (i.e., increased mobility shifts in phos-tag gel) without affecting its stability (Fig. 1C). Interestingly, GID mutation in Axin2 scaffolding protein largely abolished Axin2 phosphorylation along with protein abundance (Fig. S1A) (21). To further determine the phosphorylation status of N and its mutants, we treated it with lambda protein phosphatase (λ PPase) and subjected it to phos-tag gel. The λ PPase treatment increased mobility shift of N and point mutant of L223 or L227, while the L223/227 mutation was unaffected (Fig. 1D). Interestingly, mobility shift of N-L223/227E mutant was not affected by λ PPase treatment, suggesting that GID provides the main N phosphorylation. In a repeated experiment, we noted that N was hardly de-phosphorylated by λ PPase, while those mutants were dephosphorylated under the same experimental condition, suggesting N oligomerization by phosphorylation. To simply examine this possibility, we generated an N-EGFP fusion construct and co-transfected it with either Flag-tagged N or L223/227E mutant in 293 cells. Immunoprecipitation assay revealed that Flag-N interacts with N-EGFP, while N and L223/227E mutant interaction decreased (Fig. 1E). Immunofluorescence also supported the co-localization of N and GSK-3, especially in condensate-like structures in the cytosolic space, as compared to L223/227E mutant N (Fig. 1F). Our observations indicate that N in SARS-CoV-2 harbors GID, providing interaction with GSK-3 followed by enhanced phosphorylation of N by GSK-3.

Because the Axin-GSK-3 binding structure is well-determined (14,22), we next compared the binding site of Axin2 and N with that in wild type (wt) and mutants GSK-3. Consistently, Y216 and V267/E268 in GSK-3 were critical for Axin2 binding (Fig. S2B), and N shared those binding sites in GSK-3 (Fig. 2A), indicating that N utilizes a GSK-3 binding mode similar to that of Axin. Previously, we reported that antihelminthic niclosamide interacts with GSK-3 and inhibits Axin-GSK3 binding, resulting in attenuation of Wnt signaling and EMT in APC-mutated colorectal cancer (19). Niclosamide also inhibits viral replication of SARS-CoV-1 (23) although the molecular mechanism of action (MoA) of niclosamide on SARS-CoV is unclear. Given the similar binding modes of N and Axin to GSK-3, we next performed immunoprecipitation assay with increasing doses of niclosamide. Indeed, niclosamide inhibited Axin2 or N binding to GSK-3 in a dose-dependent manner (Fig. 2B). While N-GSK-3 interaction was maintained at 2 μM concentration of niclosamide, Axin2 binding to GSK-3 was largely abolished at the same concentration. These results suggest that N harbors stronger binding affinity to GSK-3 than does Axin2, and explain the modest therapeutic potential of niclosamide in SARS-CoV-2 in vitro (24). In turn, these results suggest the possibility that N and Axin2 compete in GSK-3 binding. To test this, we next utilized split GFP systems (25). For the split GFP assay, superfolder GFP1-10 (amino acids 1-214) and GFP11 (amino acid 214-230) fragments were fused to GSK-3 and N, respectively. Co-transfection of GSK3-GFP1-10 and N-GFP11 gave rise to green fluorescence via direct interaction between GSK-3 and N, and fluorescence was largely decreased by the overexpression of Axin2, indicating that N and Axin compete in similar binding regions of GSK-3 (Fig. 2C).

**Figure 2.**
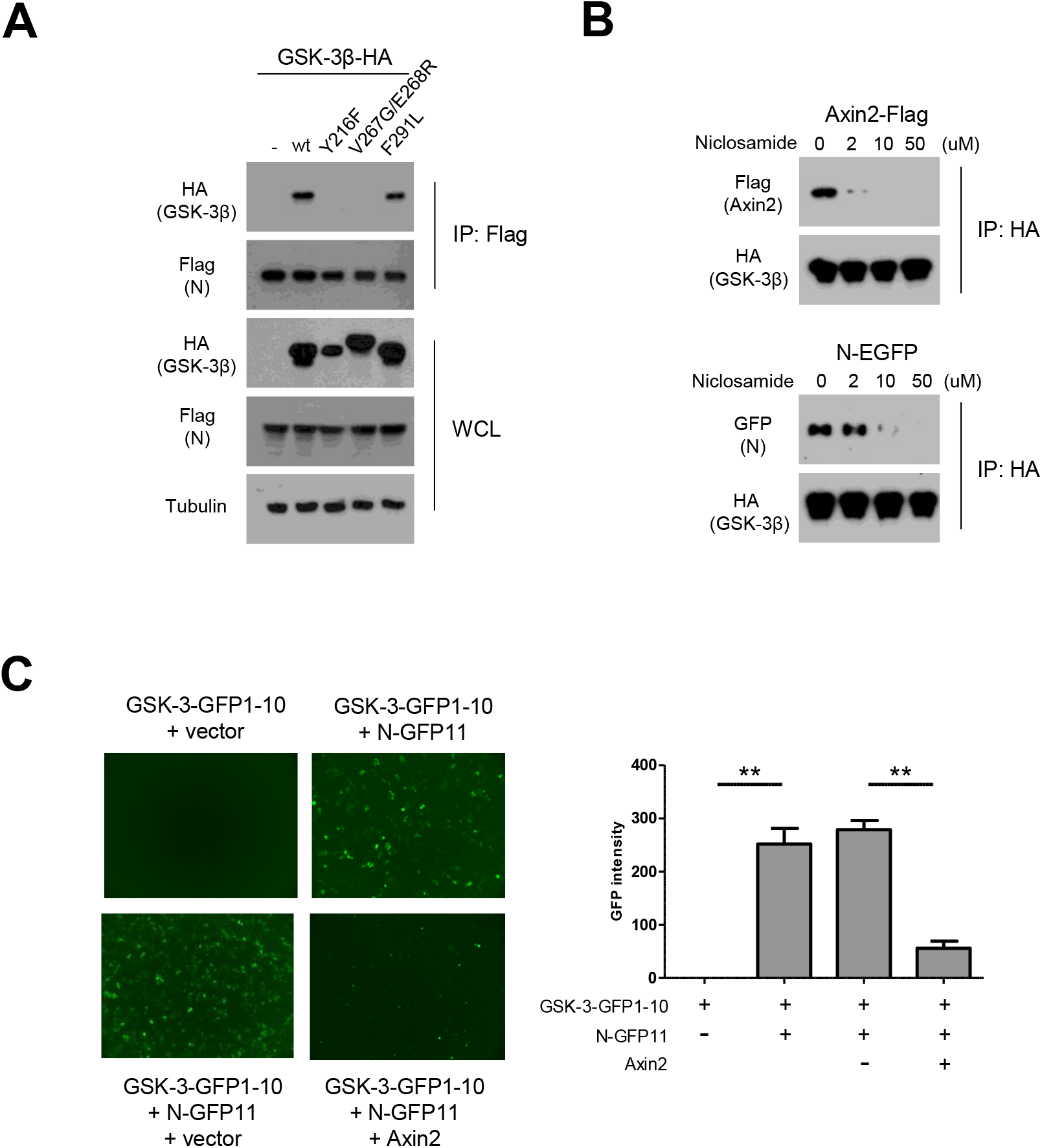
N interacts with GSK-3β similarly to Axin2. A, Flag-tagged N was co-transfected with wt (wild type) or Axin-binding mutant (Y216F, V267G/E268R, F291L) GSK-3β expression vectors in 293 cells. Following IP with anti-flag, GSK-3β binding was determined by western blot analysis. B, Flag-tagged N was co-transfected with HA-tagged wt GSK-3β in 293 cells. The lysates were subjected to IP with increasing doses of niclosamide in vitro, and dissociation of Axin2 or N was determined by western blot analysis. C, One hundred ng of each GSK-3β fused to GFP1-10 and control empty vector (upper left) or N (upper right) fused to GFP11 were co-transfected into 293 cells for split GFP analysis. Competition between N and Axin2 were determined by overexpression of 1 μg of each control empty vector (lower left) or Axin2 (lower right). Green fluorescence image was obtained from 5 random fields (x100) and the intensity was quantitatively determined using ImageJ software (right panel).

### N interaction is irrelevant to GSK-3 activity and canonical Wnt activity

Because Axin directly suppresses kinase activity of GSK-3 via GID, thus regulating various signaling pathways (11,12), we next examined the role of N in GSK-3 kinase activity and the subsequent Wnt/EMT signaling pathway. Surprisingly, overexpression of N or N-GID mutant did not affect endogenous GSK-3 kinase activity in 293 cells (Fig 3A). Furthermore, overexpression of N or N-GID mutant did not affect GSK-3 abundance or phosphorylation status on Ser9 and Y216 (Fig. 3B). As well-known downstream of β-catenin, EMT-inducer Snail abundance, TCF/LEF transcriptional, and E-cadherin promoter activity were unchanged (Fig. 3C). These results indicate that N utilizes GSK-3 for its phosphorylation but does not affect GSK-3 kinase activity and subsequent signaling pathways, probably due to rapid dissociation following phosphorylation.

**Figure 3.**
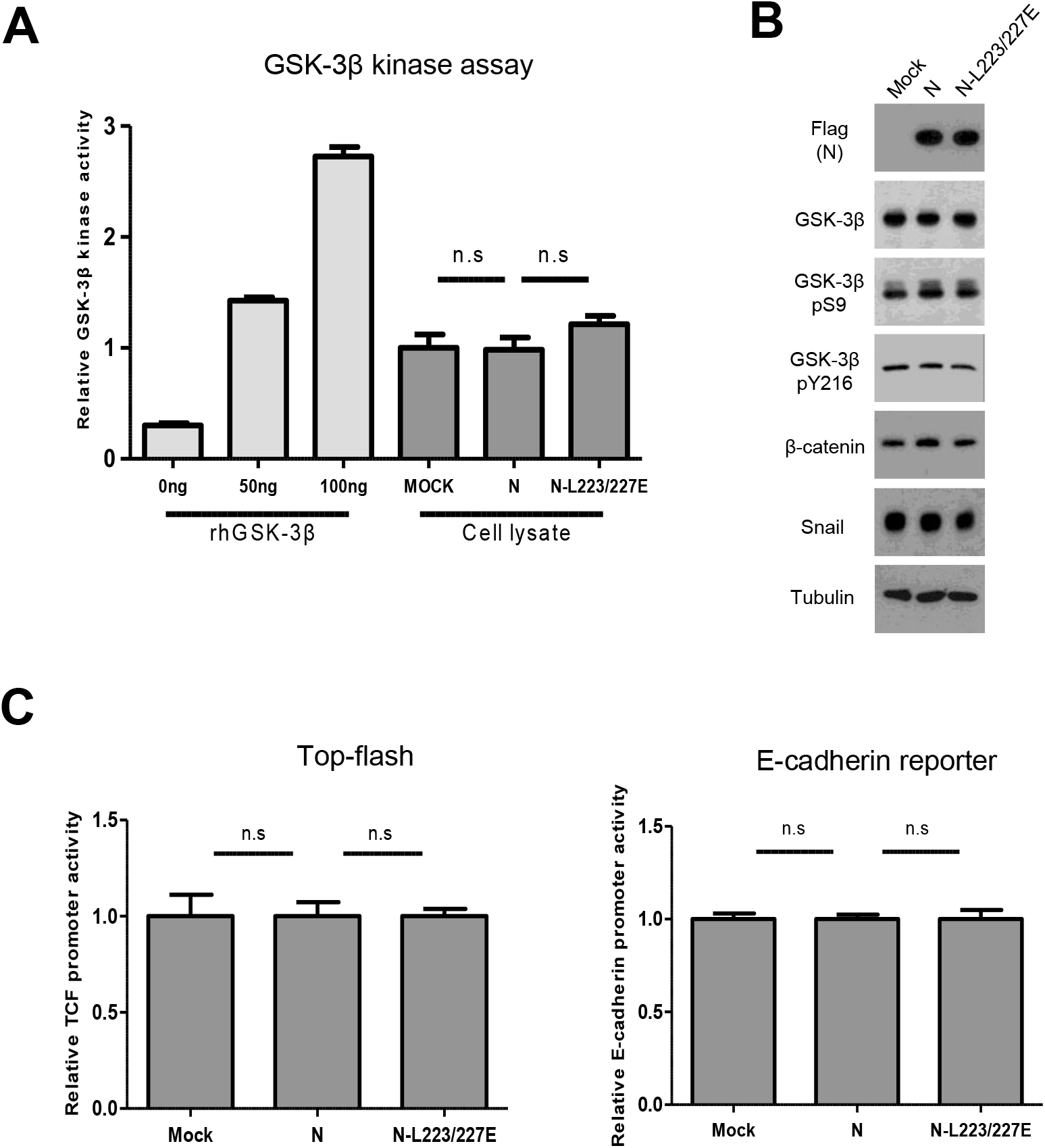
N protein does not affect endogenous GSK-3 activity and subsequent Wnt signaling and Snail abundance. A, 293 cells were transfected with N or L223/227E mutant N, and endogenous GSK-3 kinase activity was determined. rhGSK-3 in the commercial kit served as the positive control. B, N or L223/227E mutant N was overexpressed in 293 cells, and the lysates were subjected to western blot analysis to determine the phosphorylation status of GSK-3β and protein abundance of β-catenin and Snail. Tubulin served as the loading control. C, Canonical Wnt and E-cadherin transcriptional activity by N or L223/227E mutants were assessed with the Top-flash having 7x TCF/LEF binding sites (left) or Ecad(−108)-Luc, having the E-cadherin proximal promoter sequence from nt -108 to +125 (right). The reporter activity was normalized with co-transfected SV40-renilla (1 ng) with a dual-luciferase assay system.

### N-GID evolution of sarbecovirus lineage of coronavirus

While human coronaviruses HKU1, OC43, 229E, and NL63 typically cause upper respiratory infection with relatively minor symptoms, SARS-CoV and SARS-CoV-2 in the sarbecovirus lineage of beta-coronavirus replicate in the lower respiratory tract and cause severe pneumonia (26). To understand the role of GID in the evolution of coronavirus, we next analyzed GID and SR-rich phosphorylation motif among various coronaviruses (Fig. 4A). Intriguingly, we found several differences in N-GID of sarbecovirus lineage SARS-CoV-2 as compared to other coronaviruses (27). First, sarbecovirus lineage N harbors a typical GID, similar to endogenous GSK-3 binding proteins, as mentioned earlier. Second, N of sarbecovirus contained a CDK1 phosphorylation site (S206) with a conserved pSPxK/R motif that may serve to primed phosphorylation of GSK-3 (28). Third, N of sarbecovirus has a Gly-rich flexible linker between GID and SR-rich phosphorylation domain. Based on these observations, we hypothesize that CDK1 serves to primed phosphorylation of GSK-3 and that Gly-rich linker facilitates phosphorylation of N. To test this hypothesis, we generated N expression vectors with substitution either of CDK1 phosphorylation site (S206A) or GNGG linker to DEIA (N-DEIA), corresponding to sequences in HKU1 coronavirus. Then, we transfected the expression vectors and examined the protein abundance, GSK-3 binding and phosphorylation status of N. Through western blot and immunoprecipitation analysis, we found a similar protein abundance and binding capacity to GSK-3 of N and mutant N expression vectors (Fig. 4B). Interestingly, mutation of the CDK1 phosphorylation site (N-S206A) and of Gly-rich linker (N-DEIA) decreased phosphorylation status compared to ancestral N in phos-tag analysis (Fig. 4C). These results indicate that emergence of CDK1 and Gly-rich linker together with GID in sarbecovirus lineage play an important role in enhanced phosphorylation of N.

**Figure 4.**
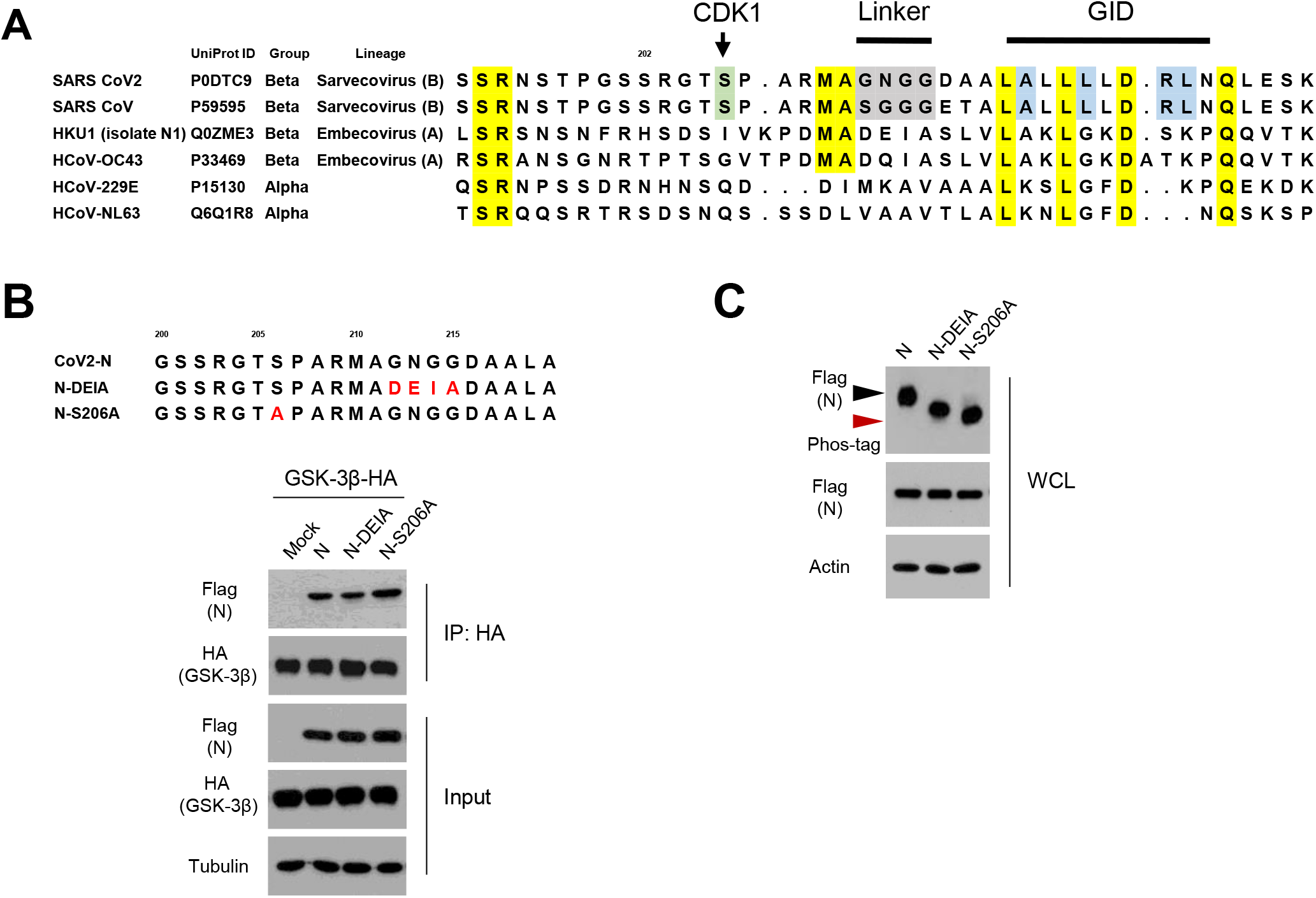
CDK1 primed phosphorylation and Gly-rich linker enhance N phosphorylation. A. N sequence between the SR-rich motif and GID in various coronavirus groups causing human respiratory infection. CDK1 phosphorylation site and Gly-rich linker are marked with green and gray, respectively. B, Sequence alignment of N expression vectors having CDK1 phosphorylation site mutant (S206A) and substitution of Gly-rich linker (from aa 212-215) to DEIA found in HKU1 coronavirus (upper). The ancestral N or N-DEIA or N-S206A were co-transfected with HA-tagged GSK-3β into 293 cells, and the lysates were subjected to IP with anti-HA and subsequent western blot analysis to determine N binding affinity. C, The ancestral N or N-DEIA or N-S206A were transfected into 293 cells and the lysates were subjected to Phos-tag gel analysis to determine N phosphorylation status. Black and red arrowheads correspond to fully phosphorylated and under-phosphorylated N on a Phos-tag gel, respectively.

### Abundance and phosphorylation status of N in other coronavirus and SARS-CoV-2 variants

Given the clinical difference between the sarbecovirus lineage of beta-coronavirus and other coronaviruses, we next compared N abundance in those coronaviruses. Transfecting N of SARS-CoV-2, 229E, OC43, and HKU1 strains, we found significantly increased N abundance in SARS-CoV-2 compared to N in other coronaviruses (Fig. 5A). Only N of 229E revealed detectable binding to GSK-3. To further determine the role of GSK-3 and GID, we next generated N-229E expression vectors having a CDK1-primed phosphorylation site, Gly-rich linker, and GID corresponding to those of sequences in N of SARS-CoV-2. Comparing N of 229E to those mutants, we found that introduction of primed phosphorylation and Gly-rich linker significantly increased phosphorylation as well protein abundance (about 2-fold) of N-229E (Fig. 5B). Notably, N-229E showed at least 7 bands in phos-tag gel analysis, indicating that N of most coronavirus is hyper-phosphorylated in SR-rich motif (29). These results indicate that N of sarbecovirus lineage has evolved to become hyper-phosphorylated with GID, CDK-1 mediated primed phosphorylation, and flexible linker between GID and SR-rich motif.

**Figure 5.**
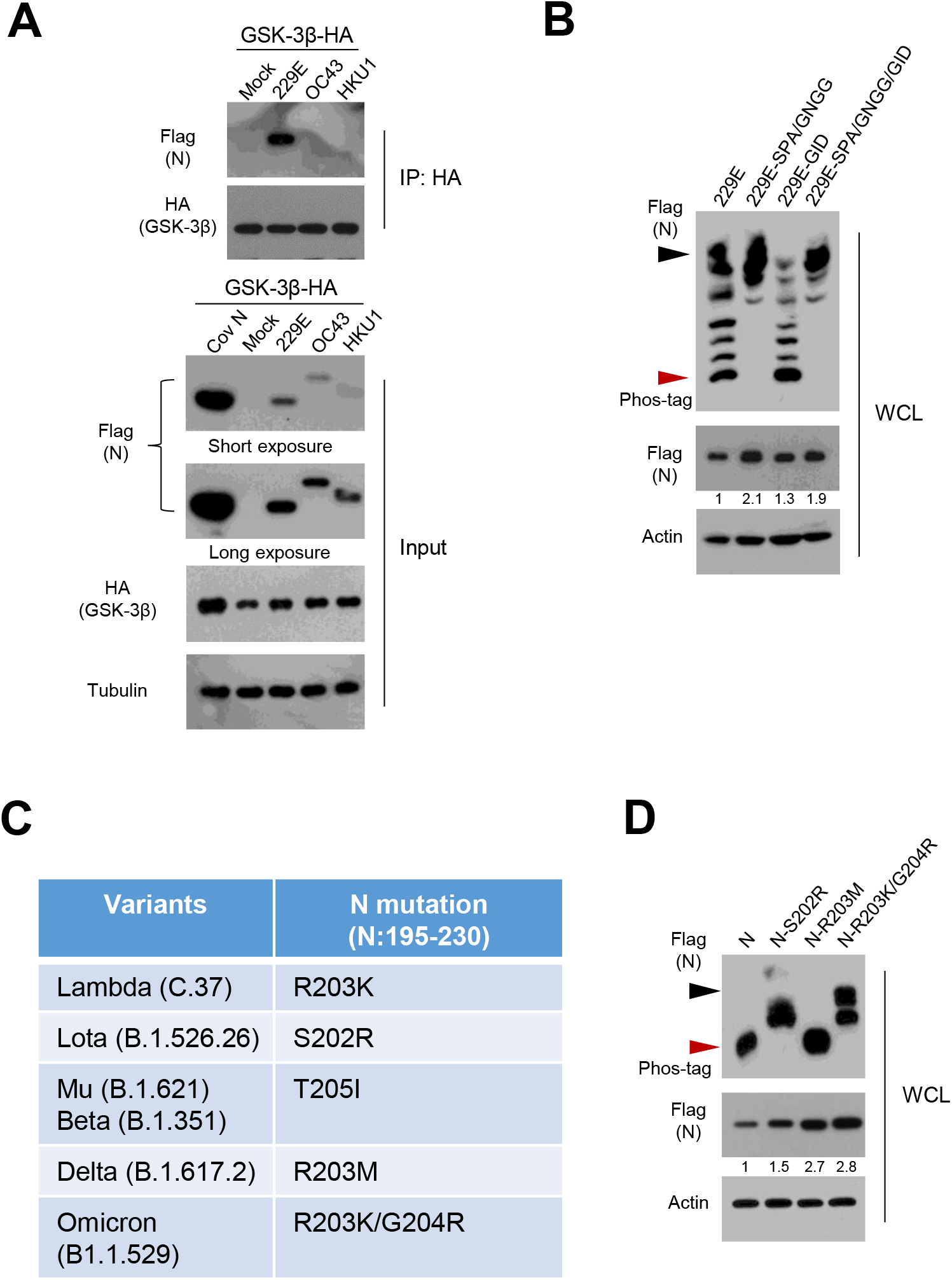
N in other coronaviruses and N variants. A, The ancestral N or N from 229E, OC43, and HKU1 were co-transfected with HA-tagged GSK-3β into 293 cells, and the lysates were subjected to IP with anti-HA and subsequent western blot analysis to determine N binding affinity. B, The N-229E or mutants having substitution of CDK1 phosphorylation site (SPAR), Gly-rich linker (GNGG), and GID with corresponding sequences of ancestral SARS-CoV-2 were transfected into 293 cells. The lysates were subjected to western blot and Phos-tag gel analysis to determine protein abundance and mobility shift by phosphorylation status of N-229, respectively. Black and red arrowheads correspond to the fully phosphorylated and under-phosphorylated N on a Phos-tag gel, respectively. C, Mutation profiles of N between SR-rich motif and GID (N195-230) in SARS-CoV-2 variants (https://cov-lineages.org/constellations). D, Ancestral N and mutants found in the variants were transfected into 293 cells. The lysates were subjected to western blot and Phos-tag gel analysis to determine protein abundance and phosphorylation status of N and mutants, respectively. Black and red arrowheads correspond to the increased phosphorylation in Omicron variant and ancestral N phosphorylation on a Phos-tag gel, respectively.

Because SARS-CoV-2 continuously evolves through mutational changes, there are many variants including Delta (B.1.617.2) and Omicron (B.1.1.529). Given our observations, we further examined the N mutation in those variants focusing on the region between SR-rich motif and GID (https://cov-lineages.org/constellations). Interestingly, we found many mutations in the variants of SARS-CoV-2, in the proximal N-terminus of the CDK1 phosphorylation site (Fig. 5C). Notably, there was no mutation among the SARS-CoV-2 variants in GID, suggesting the evolutionary fitness of GSK-3 binding in the sarbecovirus lineage. To validate the role of these mutations in the variants, we chose several mutations of N including Variants of Concern (VOC, Delta and Omicron) and generated N expression vectors having those mutations. We then overexpressed those vectors in 293 cells and examined the protein abundance and phosphorylation status of N. Interestingly, protein abundance of N in those variants was increased by the substitution of single or double amino acid(s), especially in Delta and Omicron (Fig. 5D), suggesting that N of SARS-CoV-2 is evolving to increase phosphorylation and protein abundance. The N-S202R mutant found in Lota (Variant Being Monitored, VOM) and R203K/G204R found in Omicron showed even more phosphorylation compared to ancestral N, while N found in Delta revealed a similar mobility shift, indicating that the mutational evolution provides more efficient and increased phosphorylation along with abundant N protein. Taken together, we observed that the nucleocapsid N of SARS-CoV-2 harbors hyper-phosphorylation machinery including GID, priming phosphorylation, and Gly-rich flexible linker. The evolutionary role and biological impact of N in the COVID-19 pandemic needs further elucidation, especially in BSL-3 equipped facilities.

## Discussion

GSK-3 is highly expressed endogenously in mammals and is an exceptional kinase inactivated by exogenous signaling, such as by insulin and Wnt (11,12). Priming phosphorylation by other many kinases allows strong (500-to 1,000 fold) preference of GSK-3 substrates which have multiple and serial phosphorylation sites by GSK-3, including EMT-inducer Snail and β-catenin (30,31). The importance of GSK-3 has been extensively confirmed in human diseases, including Alzheimer’s, metabolic diseases, and cancer (11,12). Notably, many viruses or bacterial genes directly interact with and utilize endogenous GSK-3 for entry, replication and latency (32). For example, latency-associated nuclear antigen (LANA) in Kaposi’s sarcoma-associated herpesvirus (KSHV) interacts with GSK-3 via GID embedded in LANA. As with Axin, the LANA-GSK-3 interaction is essential to phosphorylation of LANA and inactivation of GSK-3 (18). Cytotoxin-associated gene A (CagA) oncoprotein in *Helicobacter pylori* directly binds to GSK-3 and depletes its endogenous kinase activity, resulting in potentiation of the Snail-mediated EMT program in gastric cancer (33). Unlike LANA and CagA, N of SARS-CoV-2 does not affect endogenous GSK-3 activity and the Wnt/EMT pathway, tightly binding to GSK-3 via GID.

SARS-CoV-2 is classified as *beta-coronaviruses* genus and *Sarbecovirus* lineage (lineage B). The multifunctional nucleocapsid protein of the coronavirus family not only constitutes a key structural protein of virion but also plays a critical role in RNA binding, packaging, and other replication process (3). Upon SARS-CoV-2 infection, the N protein as well as antibodies against N appear earlier than antibodies to Spike (1,4), suggesting that N protein abundance and subsequent post-translation modification play an important role in early stage of human infection. Based on previous observations that GSK-3 phosphorylates N and its phosphorylation is critically important for viral RNA binding (6,34,35), we wondered about the role of GSK-3-mediated phosphorylation of N during SARS-CoV-2 evolution because the GSK-3 comprises a key kinase involving many endogenous biological events (11,12). We found the GID amino acid sequence in N of *Sarbecovirus* lineage consists of L/FxxxL/AxxRL. Axin binds to GSK-3 via GID, the binding being critically important for phosphorylation and protein abundance of Axin (21). However, mutation of hydrophobic residues in N-GID, which plays critical role in GSK-3 binding, largely abolished the interaction with GSK-3 without affecting N protein abundance. Our observations suggest that N, unlike GSK-3 scaffolding proteins, is highly phosphorylated by GSK-3 followed by rapid dissociation. Since the phosphorylation of N is critically important for self-oligomerization, multivalent RNA-protein complex formation, and subsequent liquid-liquid separation with RNA condensates formation (35,36), our observations suggest that GSK-3 is mainly involved in viral RNA processing within the replication transcription complex (RTC). The phosphorylation status of N protein in SARS-CoV-2 virion and existence of endogenous protein phosphatase are interesting issues raised by this study.

While SARS-CoV-2 is currently evolving increased transmissibility, the functional importance of mutations in the variants is technically challenging. Recent observation has revealed that S202R in Lota and R203M in Delta variant provide 166-fold and 51-fold higher viral production, respectively (10). In this study, we showed that pS206 priming phosphorylation by CDK1 and Gly-rich linker located between GID and SR-rich motif enhance phosphorylation of N. Given the multiple bands in N of 229E strain, N of SARS-CoV-2 may have multiple phosphorylation sites in SR-rich motif. Interestingly, emerging variants have mutations near the primed phosphorylation site. Notably, R203K/G204R, found in a highly transmissible Omicron variant, allows greater phosphorylation and protein abundance than N of ancestral and other variants. Because more than fifteen Ser/Thr residues exist in SR-rich motif, our results indicate that N of Omicron gains further phosphorylation via R203K/G204R mutation. Taken together, our observations suggest that enhanced phosphorylation and increased protein abundance of N as well as mutations of Spike may provide a selective advantage during the mutational evolution of SARS-CoV-2. Further study is needed to elucidate the importance of N phosphorylation in the emergence and evolution of SARS-CoV-2.

Previously, we observed that antiheminthic niclosamide disrupts Axin-GSK3 interaction, providing a repositioned therapeutic for colon cancer and familial adenomatosis coli (20). While niclosamide is consistently effective in disrupting Axin2-GSK-3 binding, an at least 5-fold concentration of niclosamide was required for disruption of N-GSK-3 interaction in our hands, indicating that N has a stronger interaction with GSK-3 than that of Axin2. Conversely, our observations provide an MoA of niclosamide on SARS-CoV replication and N expression, at least in part (23), with implications for the clinical trial of niclosamide for SARS-CoV-2 (37). Although GSK-3 kinase inhibitors can be a therapeutic target for SARS-CoV-2, the endogenous abundance of GSK-3 along with its diverse physiological roles largely limit the therapeutic potential of GSK-3 inhibitors for viral diseases. Thus, further study is required regarding the protein-protein interaction of N-GSK-3 as a therapeutic target for SARS-CoV-2 infection in human.

## Experimental procedures

### Expression constructs and antibodies

The 293 cells obtained from ATCC were routinely cultured in DMEM medium containing 10% FBS. The expression vector pGBW-m4134490 (plasmid number 152580) having codon optimized N of SARS-CoV, pGBW-m4134909 (plasmid number 151901) having N of human coronavirus 229E, pGBW-m4134899 (plasmid number 151902) having N of human coronavirus OC43 and pGBW-m4134901 (plasmid number 151922) having N of human coronavirus HKU1 229E were obtained from Addgene. Those N expression vectors were subcloned into pcDNA3.1 with C-terminal flag or EGFP tag. Mutant expression vectors of N in GID, linker, and CDK1 phosphorylation site were generated by a PCR-based method. The transfection was performed by Lipofectamine 2000 according to the manufacturer’s protocol (Invitrogen). Antibodies against GSK-3β (610202, BD Transduction Laboratories), pS9-GSK-3β (9323S, Cell Signaling), pY216-GSK-3β (612312, BD Transduction Laboratories), Flag (F-3165, Sigma), Snail (L70G2, Cell Signaling), β-catenin (610154, BD Transduction Laboratories), and Tubulin (LF-PA0146, AbFrontier) were obtained from the commercial vendors. Endogenous GSK-3β kinase activity was determined by commercial kit (V1991, Promega).

### Western blot and Phos-tag gel analysis

For the western blot analyses, protein extracts were prepared in Triton X-100 lysis buffer. Phosphorylation status of N protein was determined by anti-flag antibody and mobility shift on a Phos-tag gel (Wako) as described previously (38).

### Immunoprecipitation and immunofluorescence

For immunoprecipitation analysis, whole cell Triton X-100 lysates were incubated with Flag-M2 agarose (Sigma) and washed with lysis buffer three times. The recovered proteins were resolved by SDS-PAGE and subjected to immunoblot analysis. For immunofluorescence study, the cells were washed twice with ice-cold PBS and incubated for 15 min at room temperature with 3% formaldehyde in PBS. The cells were permeabilized with 0.5% Triton X-100 for 5 min and then blocked for 1 h in PBS containing 3% bovine serum albumin followed by incubation with primary antibody over night at 4 °C. Cells were then washed three times with PBS containing 0.1% Tween 20 followed by incubation with anti-mouse-Alexa Fluor-488 (for green) or anti-rabbit-Alexa Fluor-594 (for red) secondary antibody. Cellular fluorescence was monitored using confocal microscopy (Zeiss).

### TCF/LEF and E-cadherin reporter assay

For TCF/LEF and E-cadherin reporter assay, the cells were transfected with 50 ng of the Super-Top or E-cad(−108)-Luc reporter vector (30,39) and 1 ng of pSV40-Renilla expression vector in combination with N or N-mutant as indicated. Luciferase and *renilla* activities were measured using the dual-luciferase reporter system kit (Promega), and the luciferase activity was normalized with *renilla* activity. The results are expressed as the averages of the ratios of the reporter activities from triplicate experiments.

### Split superfolder GFP assay

For split superfolder GFP assay, the GFP1-10 and GFP11 constructs were kindly provided by Professor Hye Yoon Park at Seoul National University. The GFP1-10 fragment was fused into N-terminus of GSK-3 and GFP11 was fused into C-terminus of N. The 293 cells were co-transfected with split GFP vectors, and fluorescence intensity was determined by 5 random areas with the same exposure, followed by ImageJ analysis.

### Protein sequences of coronaviruses and SARS-CoV-2 variants

Amino acid sequences of N for SARS-CoV-2 (P0DTC9), SARS-CoV (P59595), HKU1 (Q0ZME3), HCoV-OC43 (P33469), HCoV-229E (PP15130), HCoV-NL63 (Q6Q1R8) were obtained from UniProt (https://www.uniprot.org/). Mutational information on N protein in SARS-CoV-2 variants was obtained from PANGO Lineages (https://cov-lineages.org/constellations)

### Statistics

Statistical analysis for reporter, GSK-3 kinase, and split GFP assay was performed with two-tailed Student’s *t*-test; data are expressed as means and s.d. The double asterisks denote *p* < 0.01, one asterisk denoting *p* < 0.05.

## ACKOWLEDGMENTS

We thank E. Tunkle for preparation of the manuscript and J. Choi at Dongduk University College of Pharmacy for technical assistance. This work was supported by grants from the National Research Foundation of Korea (NFR-2017R1A2B3002241, NRF-2019R1A2C2084535, NRF-2021R1A2C3003496) funded by the Korean government (MSIP), a grant from the National Research Foundation of Korea (NRF-2020R1I1A1A01072977) funded by the Korea government (MOE).

## AUTHOR CONTRIBUTIONS

J.S.Y. performed all experiments; H.S., S.Y.C., J.E.L., C-H.J., S.H.S., S.K., and E.S.C. supported experiments; N.H.K. and K.H.H. performed split GFP assay; N.H.K., H.S.K., and J.I.Y. planned all experiments, analyzed the data, and wrote the manuscript.

## FIGURE LEGENDS

**Fig. S1.**
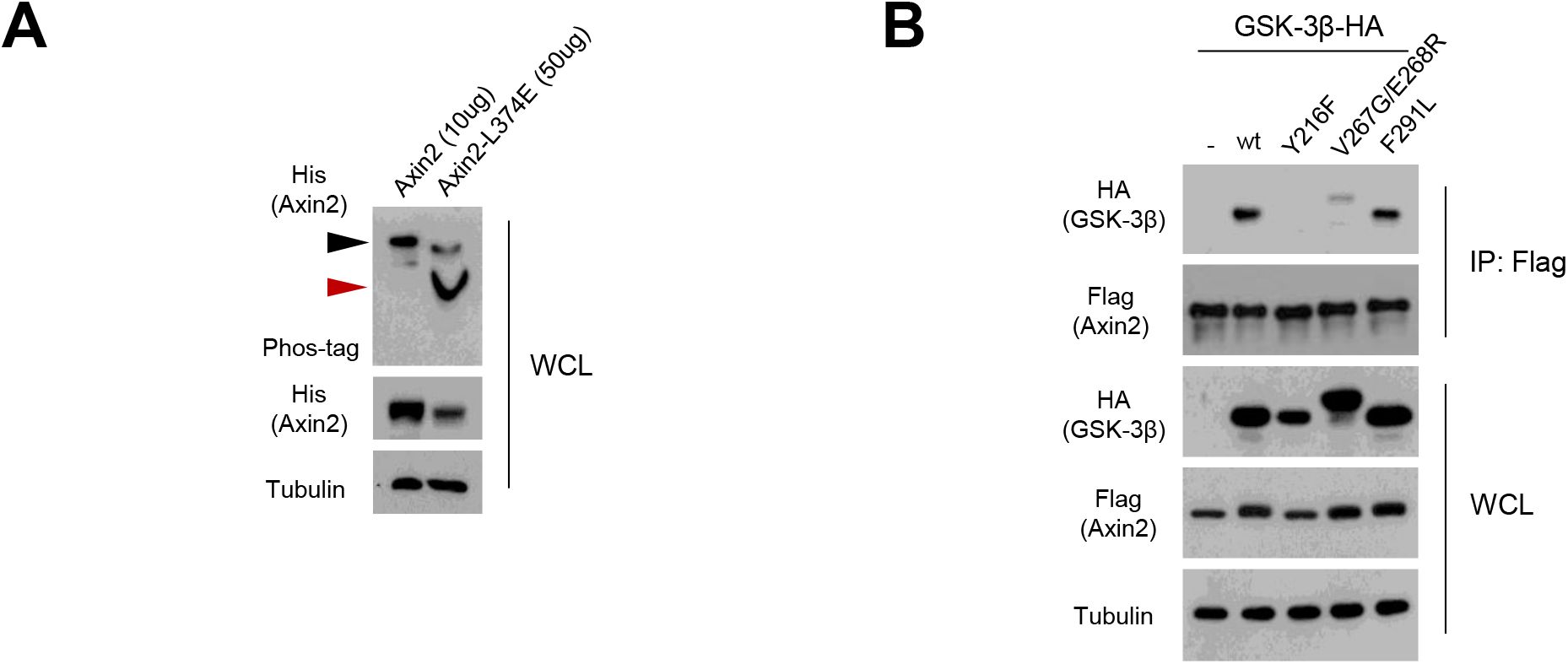
A. Axin2 phosphorylation and its stability were dependent on GSK-3β binding. Flag-tagged Axin2 or mutants harboring Lue to Gly substitution in GSK-3 β binding site (L374E) were co-transfected with HA-tagged GSK-3β in 293 cells. The cell lysates were subjected to western blot and Phos-tag gel analysis to determine protein abundance and phosphorylation status, respectively. For Phos-tag gel analysis, 10 μg of wt lysate and 50 μg of Axin2-L374E were loaded to compare phosphorylation status. **Fig. S1B. Axin2 share GSK-3β site with N**. Flag-tagged Axin2 was co-transfected with wt (wild type) or Axin-binding mutants (Y216F, V267G/E268R, F291L) GSK-3β expression vectors in 293 cells. Following IP with anti-flag, GSK-3β binding was determined using western blot analysis.

## REFERENCES

1. Shan, D., Johnson, J. M., Fernandes, S. C., Suib, H., Hwang, S., Wuelfing, D., Mendes, M., Holdridge, M., Burke, E. M., Beauregard, K., Zhang, Y., Cleary, M., Xu, S., Yao, X., Patel, P. P., Plavina, T., Wilson, D. H., Chang, L., Kaiser, K. M., Nattermann, J., Schmidt, S. V., Latz, E., Hrusovsky, K., Mattoon, D., and Ball, A. J. (2021) N-protein presents early in blood, dried blood and saliva during asymptomatic and symptomatic SARS-CoV-2 infection. Nat Commun 12, 1931

2. de Haan, C. A., and Rottier, P. J. (2005) Molecular interactions in the assembly of coronaviruses. Adv Virus Res 64, 165–230

3. Chang, C. K., Hou, M. H., Chang, C. F., Hsiao, C. D., and Huang, T. H. (2014) The SARS coronavirus nucleocapsid protein--forms and functions. Antiviral Res 103, 39–50

4. Tan, Y. J., Goh, P. Y., Fielding, B. C., Shen, S., Chou, C. F., Fu, J. L., Leong, H. N., Leo, Y. S., Ooi, E. E., Ling, A. E., Lim, S. G., and Hong, W. (2004) Profiles of antibody responses against severe acute respiratory syndrome coronavirus recombinant proteins and their potential use as diagnostic markers. Clin Diagn Lab Immunol 11, 362–371

5. Krutikov, M., Palmer, T., Tut, G., Fuller, C., Shrotri, M., Williams, H., Davies, D., Irwin-Singer, A., Robson, J., Hayward, A., Moss, P., Copas, A., and Shallcross, L. (2021) Incidence of SARS-CoV-2 infection according to baseline antibody status in staff and residents of 100 long-term care facilities (VIVALDI): a prospective cohort study. Lancet Healthy Longev 2, e362–e370

6. Wu, C. H., Yeh, S. H., Tsay, Y. G., Shieh, Y. H., Kao, C. L., Chen, Y. S., Wang, S. H., Kuo, T. J., Chen, D. S., and Chen, P. J. (2009) Glycogen synthase kinase-3 regulates the phosphorylation of severe acute respiratory syndrome coronavirus nucleocapsid protein and viral replication. J Biol Chem 284, 5229–5239

7. Peng, T. Y., Lee, K. R., and Tarn, W. Y. (2008) Phosphorylation of the arginine/serine dipeptide-rich motif of the severe acute respiratory syndrome coronavirus nucleocapsid protein modulates its multimerization, translation inhibitory activity and cellular localization. Febs j 275, 4152–4163

8. Wu, C. H., Chen, P. J., and Yeh, S. H. (2014) Nucleocapsid phosphorylation and RNA helicase DDX1 recruitment enables coronavirus transition from discontinuous to continuous transcription. Cell Host Microbe 16, 462–472

9. Liu, X., Verma, A., Garcia, G., Jr., Ramage, H., Lucas, A., Myers, R. L., Michaelson, J. J., Coryell, W., Kumar, A., Charney, A. W., Kazanietz, M. G., Rader, D. J., Ritchie, M. D., Berrettini, W. H., Schultz, D. C., Cherry, S., Damoiseaux, R., Arumugaswami, V., and Klein, P. S. (2021) Targeting the coronavirus nucleocapsid protein through GSK-3 inhibition. Proc Natl Acad Sci U S A 118

10. Syed, A. M., Taha, T. Y., Tabata, T., Chen, I. P., Ciling, A., Khalid, M. M., Sreekumar, B., Chen, P. Y., Hayashi, J. M., Soczek, K. M., Ott, M., and Doudna, J. A. (2021) Rapid assessment of SARS-CoV-2-evolved variants using virus-like particles. Science 374, 1626–1632

11. Cohen, P., and Frame, S. (2001) The renaissance of GSK3. Nat Rev Mol Cell Biol 2, 769–776

12. Doble, B. W., and Woodgett, J. R. (2003) GSK-3: tricks of the trade for a multi-tasking kinase. J Cell Sci 116, 1175–1186

13. Hedgepeth, C. M., Deardorff, M. A., Rankin, K., and Klein, P. S. (1999) Regulation of glycogen synthase kinase 3beta and downstream Wnt signaling by axin. Mol Cell Biol 19, 7147–7157

14. Dajani, R., Fraser, E., Roe, S. M., Yeo, M., Good, V. M., Thompson, V., Dale, T. C., and Pearl, L. H. (2003) Structural basis for recruitment of glycogen synthase kinase 3beta to the axin-APC scaffold complex. Embo j 22, 494–501

15. Howng, S. L., Hwang, C. C., Hsu, C. Y., Hsu, M. Y., Teng, C. Y., Chou, C. H., Lee, M. F., Wu, C. H., Chiou, S. J., Lieu, A. S., Loh, J. K., Yang, C. N., Lin, C. S., and Hong, Y. R. (2010) Involvement of the residues of GSKIP, AxinGID, and FRATtide in their binding with GSK3beta to unravel a novel C-terminal scaffold-binding region. Mol Cell Biochem 339, 23–33

16. Wang, H. Y., Juo, L. I., Lin, Y. T., Hsiao, M., Lin, J. T., Tsai, C. H., Tzeng, Y. H., Chuang, Y. C., Chang, N. S., Yang, C. N., and Lu, P. J. (2012) WW domain-containing oxidoreductase promotes neuronal differentiation via negative regulation of glycogen synthase kinase 3β. Cell Death Differ 19, 1049–1059

17. Yook, J. I., Li, X. Y., Ota, I., Hu, C., Kim, H. S., Kim, N. H., Cha, S. Y., Ryu, J. K., Choi, Y. J., Kim, J., Fearon, E. R., and Weiss, S. J. (2006) A Wnt-Axin2-GSK3beta cascade regulates Snail1 activity in breast cancer cells. Nat Cell Biol 8, 1398–1406

18. Fujimuro, M., Liu, J., Zhu, J., Yokosawa, H., and Hayward, S. D. (2005) Regulation of the interaction between glycogen synthase kinase 3 and the Kaposi’s sarcoma-associated herpesvirus latency-associated nuclear antigen. J Virol 79, 10429–10441

19. Ahn, S. Y., Kim, N. H., Lee, K., Cha, Y. H., Yang, J. H., Cha, S. Y., Cho, E. S., Lee, Y., Cha, J. S., Cho, H. S., Jeon, Y., Yuk, Y. S., Cho, S., No, K. T., Kim, H. S., Lee, H., Choi, J., and Yook, J. I. (2017) Niclosamide is a potential therapeutic for familial adenomatosis polyposis by disrupting Axin-GSK3 interaction. Oncotarget 8, 31842–31855

20. Ahn, S. Y., Yang, J. H., Kim, N. H., Lee, K., Cha, Y. H., Yun, J. S., Kang, H. E., Lee, Y., Choi, J., Kim, H. S., and Yook, J. I. (2017) Anti-helminthic niclosamide inhibits Ras-driven oncogenic transformation via activation of GSK-3. Oncotarget 8, 31856–31863

21. Yamamoto, H., Kishida, S., Kishida, M., Ikeda, S., Takada, S., and Kikuchi, A. (1999) Phosphorylation of axin, a Wnt signal negative regulator, by glycogen synthase kinase-3beta regulates its stability. J Biol Chem 274, 10681–10684

22. Fraser, E., Young, N., Dajani, R., Franca-Koh, J., Ryves, J., Williams, R. S., Yeo, M., Webster, M. T., Richardson, C., Smalley, M. J., Pearl, L. H., Harwood, A., and Dale, T. C. (2002) Identification of the Axin and Frat binding region of glycogen synthase kinase-3. J Biol Chem 277, 2176–2185

23. Wu, C. J., Jan, J. T., Chen, C. M., Hsieh, H. P., Hwang, D. R., Liu, H. W., Liu, C. Y., Huang, H. W., Chen, S. C., Hong, C. F., Lin, R. K., Chao, Y. S., and Hsu, J. T. (2004) Inhibition of severe acute respiratory syndrome coronavirus replication by niclosamide. Antimicrob Agents Chemother 48, 2693–2696

24. Ko, M., Jeon, S., Ryu, W. S., and Kim, S. (2021) Comparative analysis of antiviral efficacy of FDA-approved drugs against SARS-CoV-2 in human lung cells. J Med Virol 93, 1403–1408

25. Cabantous, S., Terwilliger, T. C., and Waldo, G. S. (2005) Protein tagging and detection with engineered self-assembling fragments of green fluorescent protein. Nat Biotechnol 23, 102–107

26. Tay, M. Z., Poh, C. M., Rénia, L., MacAry, P. A., and Ng, L. F. P. (2020) The trinity of COVID-19: immunity, inflammation and intervention. Nat Rev Immunol 20, 363–374

27. Tabibzadeh, A., Esghaei, M., Soltani, S., Yousefi, P., Taherizadeh, M., Safarnezhad Tameshkel, F., Golahdooz, M., Panahi, M., Ajdarkosh, H., Zamani, F., and Karbalaie Niya, M. H. (2021) Evolutionary study of COVID-19, severe acute respiratory syndrome coronavirus 2 (SARS-CoV-2) as an emerging coronavirus: Phylogenetic analysis and literature review. Vet Med Sci 7, 559–571

28. Surjit, M., Liu, B., Chow, V. T., and Lal, S. K. (2006) The nucleocapsid protein of severe acute respiratory syndrome-coronavirus inhibits the activity of cyclin-cyclin-dependent kinase complex and blocks S phase progression in mammalian cells. J Biol Chem 281, 10669–10681

29. Lin, L., Shao, J., Sun, M., Liu, J., Xu, G., Zhang, X., Xu, N., Wang, R., and Liu, S. (2007) Identification of phosphorylation sites in the nucleocapsid protein (N protein) of SARS-coronavirus. Int J Mass Spectrom 268, 296–303

30. Yook, J. I., Li, X. Y., Ota, I., Fearon, E. R., and Weiss, S. J. (2005) Wnt-dependent regulation of the E-cadherin repressor snail. J Biol Chem 280, 11740–11748

31. Kaidanovich-Beilin, O., and Woodgett, J. R. (2011) GSK-3: Functional Insights from Cell Biology and Animal Models. Front Mol Neurosci 4, 40

32. Alfhili, M. A., Alsughayyir, J., McCubrey, J. A., and Akula, S. M. (2020) GSK-3-associated signaling is crucial to virus infection of cells. Biochim Biophys Acta Mol Cell Res 1867, 118767

33. Lee, D. G., Kim, H. S., Lee, Y. S., Kim, S., Cha, S. Y., Ota, I., Kim, N. H., Cha, Y. H., Yang, D. H., Lee, Y., Park, G. J., Yook, J. I., and Lee, Y. C. (2014) Helicobacter pylori CagA promotes Snail-mediated epithelial-mesenchymal transition by reducing GSK-3 activity. Nat Commun 5, 4423

34. Chen, H., Gill, A., Dove, B. K., Emmett, S. R., Kemp, C. F., Ritchie, M. A., Dee, M., and Hiscox, J. A. (2005) Mass spectroscopic characterization of the coronavirus infectious bronchitis virus nucleoprotein and elucidation of the role of phosphorylation in RNA binding by using surface plasmon resonance. J Virol 79, 1164–1179

35. Lu, S., Ye, Q., Singh, D., Cao, Y., Diedrich, J. K., Yates, J. R., 3rd, Villa, E., Cleveland, D. W., and Corbett, K. D. (2021) The SARS-CoV-2 nucleocapsid phosphoprotein forms mutually exclusive condensates with RNA and the membrane-associated M protein. Nat Commun 12, 502

36. Carlson, C. R., Asfaha, J. B., Ghent, C. M., Howard, C. J., Hartooni, N., and Morgan, D. O. (2020) Phosphorylation modulates liquid-liquid phase separation of the SARS-CoV-2 N protein. bioRxiv

37. Al-Kuraishy, H. M., Al-Gareeb, A. I., Alzahrani, K. J., Alexiou, A., and Batiha, G. E. (2021) Niclosamide for Covid-19: bridging the gap. Mol Biol Rep 48, 8195–8202

38. Lee, Y., Kim, N. H., Cho, E. S., Yang, J. H., Cha, Y. H., Kang, H. E., Yun, J. S., Cho, S. B., Lee, S. H., Paclikova, P., Radaszkiewicz, T. W., Bryja, V., Kang, C. G., Yuk, Y. S., Cha, S. Y., Kim, S. Y., Kim, H. S., and Yook, J. I. (2018) Dishevelled has a YAP nuclear export function in a tumor suppressor context-dependent manner. Nat Commun 9, 2301

39. Kim, N. H., Kim, H. S., Li, X. Y., Lee, I., Choi, H. S., Kang, S. E., Cha, S. Y., Ryu, J. K., Yoon, D., Fearon, E. R., Rowe, R. G., Lee, S., Maher, C. A., Weiss, S. J., and Yook, J. I. (2011) A p53/miRNA-34 axis regulates Snail1-dependent cancer cell epithelial-mesenchymal transition. J Cell Biol 195, 417–433

